# Whole genome sequence-based haplotypes reveal single origin of the sickle allele during the Holocene Wet Phase

**DOI:** 10.1101/187419

**Authors:** Daniel Shriner, Charles N. Rotimi

## Abstract

Five classical designations of sickle haplotypes are based on the presence/absence of restriction sites and named after ethnic groups or geographic regions from which patients originated. Each haplotype is thought to represent an independent occurrence of the sickle mutation. We investigated the origins of the sickle mutation using whole genome sequence data. We identified 156 carriers from the 1000 Genomes Project, the African Genome Variation Project, and Qatar. We defined a new haplotypic classification using 27 polymorphisms in linkage disequilibrium with rs334. Network analysis revealed a common haplotype that differed from the ancestral haplotype only by the derived sickle mutation at rs334 and correlated collectively with the Central African Republic/Bantu, Cameroon, and Arabian/Indian designations. Other haplotypes were derived from this haplotype and fell into two clusters, one comprised of haplotypes correlated with the Senegal designation and the other comprised of haplotypes correlated with both the Benin and Senegal designations. The near-exclusive presence of the original sickle haplotype in the Central African Republic, Kenya, Uganda, and South Africa is consistent with this haplotype predating the Bantu Expansion. Modeling of balancing selection indicated that the heterozygote advantage was 15.2%, an equilibrium frequency of 12.0% was reached after 87 generations, and the selective environment predated the mutation. The posterior distribution of the ancestral recombination graph yielded an age of the sickle mutation of 259 generations, corresponding to 7,300 years and the Holocene Wet Phase. These results clarify the origin of the sickle allele and improve and simplify the classification of sickle haplotypes.

## INTRODUCTION

Several hereditary variants in the hemoglobin genes afford protection against malaria. Many such variants are thought to have evolved in the last 10,000 years.^1; 2^ In particular, the sickle allele β^S^ in the beta globin gene *HBB is* a polymorphism under balancing selection due to recessive lethality and heterozygote advantage. The chromosomal background of the β^S^ allele has been classified based on the presence or absence of a set of seven canonical restriction sites, 5’ ε HincII – Gγ1 HindIII – Aγ1 HindIII – Ψβ HincII – 3’ Ψβ HincII – βAvaII – 3’ β BamHI, yielding five haplotypes named after ethno-linguistic groups or geographic regions, *i.e*., Arabian/Indian, Benin, Cameroon, Central African Republic/Bantu, and Senegal, as well as a sixth category for “atypical” haplotypes.^3-9^

Whether the β^S^ allele has a recent or old origin has been debated since the development of restriction fragment length polymorphism data.^10; 11^ According to the multicentric model, the origin of the β^S^ allele is recent, within the last few thousand years, and each haplotype represents an independent occurrence of the same exact mutation in the corresponding geographic region. ^4^; ^12-14^ In contrast, according to the unicentric model, the origin of the β^S^ allele is old, anywhere from tens to hundreds of thousands of years, and the mutation occurred once.^15-18^ Suggested places of origin include Equatorial Africa^19^ and the Middle East.^20; 21^ kb recombination hotspot exists 1 kb upstream of *HBB*.^22^ Consequently, recombination and gene conversion, rather than *de novo* mutation, have generated several haplotypes.^3; 23; 24^

We investigated the origins of the sickle allele using whole genome sequence data from the 1000 Genomes Project, the African Genome Variation Project, and Qatar. We identified a total of 156 sickle carriers. Using phased sequence data, we established a new haplotypic classification. We then used a combination of forward time simulation, phylogenetic network analysis, and coalescent analysis to infer a single origin of the sickle allele approximately 7,300 years ago, during the Holocene Wet Phase or Green Sahara.

## MATERIALS AND METHODS

### Ethics Statement

This project was excluded from IRB review by the Office of Human Subjects Research Protections, National Institutes of Health (OHSRP ID# 17-NHGRI-00282).

### Sequence Data

We retrieved whole genome sequence data from the 1000 Genomes Project,^25^ the African Genome Variation Project,^26^ and Qatar.^27^ Haplotypes were delimited by positions ±500 kb of rs334 and pairwise *r*^2^ ≥ 0.2 with rs334. All data were processed using VCFtools version 0.1.14.^28^

### African Ancestry

Y chromosome haplogroups were called using YFitter.^29^ Mitochondrial DNA haplogroups were called using HaploFind.^30^ Autosomal ancestry was analyzed using projection analysis in ADMIXTURE version 1.3,^31^ using a global reference panel of 21 global ancestries.^32^ To determine standard errors for the proportions of ancestral components for each individual, we reran ADMIXTURE with the addition of 200 bootstrap replicates. Accounting for both within and between individual variances, we calculated the proportions for average ancestry using inverse variance weights. We then calculated 95% confidence intervals for each ancestry and individual, zeroed out any average proportions for which the 95% confidence intervals included 0, and renormalized the remaining averages to sum to 1.

### Balancing Selection

Let the genotype frequencies of the sickle homozygote, heterozygote, and wild type homozygote be *p*^2^, 2*pq*, and *q*^2^, respectively.^2^ Let the corresponding relative fitnesses be 0, 2 *p* 1 + *s*, and 1, respectively. Then, at equilibrium, 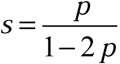. For each of the five continental African samples in the 1000 Genomes Project Phase 3 release version 5a, we estimated the effective population size *N*_*e*_ based on the heterozygosities of all single nucleotide polymorphisms (*i.e*., diallelic, triallelic, and quadrallelic), assuming a mutation rate of 0.97×10^−8^ mutations/site/generation.^33^ We then took the harmonic mean of the five *N*_*e*_ estimates. We simulated 1,000 generations under a combination of random genetic drift and balancing selection, assuming one initial copy of the mutant allele. We repeated this process 1,000 times.

### Phylogenetic network analysis

We used SplitsTree version 4.13.1 to perform split decomposition analysis of haplotypes.^34^

### Inferring the ancestral recombination graph

We used ARGweaver to infer the ancestral recombination graph.^35^ ARGweaver is based on the standard coalescent model and is sensitive to balancing selection, such that regions under balancing selection have older times to the most recent common ancestor than comparable neutral regions. We set the effective population size to the value described in the *Balancing Selection* subsection, the mutation rate to 0.97×10^−8^ mutations/generation/site,^33^ and the recombination rate to 1.5×10^−8^ recombinations/generation/site. We also investigated larger recombination rates of 1.7×10^−8^, 2.0×10^−8^, and 2.0×10^−7^ recombinations/generation/site. We used the functions heidel.diag and geweke.diag in the coda library of R, version 3.2.3, to assess convergence diagnostics based on the posterior distribution of the number of recombination events ^36^ (Figure S1). To convert generations into years, we assumed a generation interval of 28 years.^37; 38^

## RESULTS

### Molecular Mapping of Restriction Sites

We mapped 15 restriction sites, including the 7 canonical sites, to the reference human genome sequence. We identified 12 known single nucleotide polymorphisms that predict the presence or absence of 10 of these sites. Of the canonical sites, we predict 5’ ε HincII using rs3834466, Gγ1 HindIII using rs2070972, Aγ1 HindIII using rs28440105, and 3’ Ψβ HincII using rs968857 (Table 1); similar results were obtained using rs3834466, rs28440105, rs10128556, and rs968857.^39^ Correlation (measured via *r*^2^) between these variants and rs334 is weak to nonexistent (Table 1).

**Table 1.** Molecular characterization of the classical sickle designations.

| Site | Sequence <sup>a</sup> | Range | rsID | Position (hg19) | Senegal | Benin | CAR/<br>Bantu | Cameroon | Arabian/<br>Indian | r<sup>2</sup><sup>b</sup> | D'<sup>b</sup> | Ancestral | Status | Derived | Status |
| --- | --- | --- | --- | --- | --- | --- | --- | --- | --- | --- | --- | --- | --- | --- | --- |
| 3' ψβ HincII | G**T**TGAC | 5260457-5260462 | rs968857 | 5260458 | + | + | - | + | + | 0.000 | -0.104 | T | + | C | - |
| Aγ1 HindIII | **A**AGCTT | 5269799-5269804 | rs28440105 | 5269799 | - | - | - | + | - | 0.016 | 0.930 | C | - | A | + |
| Gγ1 HindIII | **A**AGCTT | 5274717-5274722 | rs2070972 | 5274717 | + | - | + | + | + | 0.003 | 0.094 | C | - | A | + |
| 5' ε HincII | G**T**TGAC | 5291563-5291567 | rs3834466 | 5291563-5291564 | - | - | - | - | + | 0.020 | -0.853 | G | - | GT | + |

<sup>a</sup> Red indicates the polymorphic position.
<sup>b</sup> Pairwise linkage disequilibrium values are shown with respect to rs334.

### Distributions of β^S^ and the classical haplotypes

In the 1000 Genomes Project, we identified 137 sickle carriers and 0 sickle homozygotes; we predicted the classical haplotypes for all 137 carriers (Table 2). The average sickle allele frequency was 12.0% and did not statistically differ among the five continental African samples ( 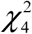 =1.48, *p*= 0.830). The distribution of matrilines comprised 1 A2, 2 B2, 1 J2, 8 L0, 24 L1, 43 L2, 49 L3, 3 L4, 2 L5, 1 T2, and 3 U6 haplogroups. The distribution of patrilines comprised 2 A1a, 5 E1a, 54 E1b1a, 1 E1b1b, 2 E2b, 1 G2a, 1 I2a, and 2 R1b. Of the 54 E1b1a, 29 were E1b1a1a1f and 23 were E1b1a1a1g.

**Table 2.** Distribution of classical sickle designations.

| Sample <sup>a</sup> | Arabian/Indian | Benin | Cameroon | CAR/Bantu | Senegal | Atypical |
| --- | --- | --- | --- | --- | --- | --- |
| ACB | 0 | 4 | 0 | 2 | 3 | 0 |
| ASW | 0 | 1 | 1 | 0 | 0 | 0 |
| CLM | 0 | 0 | 0 | 1 | 0 | 1 |
| ESN | 0 | 18 | 1 | 0 | 5 | 0 |
| GWD | 0 | 2 | 0 | 0 | 24 | 0 |
| LWK | 1 | 0 | 0 | 19 | 0 | 0 |
| MSL | 0 | 3 | 0 | 0 | 17 | 1 |
| PUR | 0 | 0 | 0 | 1 | 2 | 0 |
| YRI | 0 | 19 | 0 | 0 | 10 | 1 |
| Baganda | 0 | 0 | 0 | 14 | 0 | 0 |
| Zulu | 0 | 0 | 0 | 1 | 0 | 0 |

<sup>a</sup> The population codes are: ACB, “African Caribbean in Barbados”; ASW, “People with African Ancestry in Southwest USA”; CLM, “Colombians in Medellín, Colombia”; ESN, “Esan in Nigeria”; GWD, “Gambian in Western Division, Mandinka”; LWK, “Luhya in Webuye, Kenya”; MSL, “Mende in Sierra Leone”; PUR, “Puerto Ricans in Puerto Rico”; and YRI, “Yoruba in Ibadan, Nigeria”.

In the African Genome Variation Project, we identified 14 sickle carriers in the Baganda and 1 sickle carrier in the Zulu. We predicted that all 15 of these individuals carried the Central African Republic/Bantu haplotype (Table 2). In the Qatar sample, we identified 4 sickle carriers, all with insufficient information to predict the haplotypes.

### New classification of haplotypes based on linkage disequilibrium

We defined haplotypes centered on rs334 in the 504 continental Africans from the 1000 Genomes Project. First, we extracted 18,402 sites within 500 kb of rs334 with any non-reference allele count of 1. Then, we recorded pairwise LD for phased, diallelic sites (Table S1). The largest value of *r*^2^ with rs334 was 0.407 and there were 27 sites with *r*^2^ ≥ 0.2. Based on rs334 and these 27 sites, we observed 62 haplotypes, of which 18 carried the sickle allele at rs334; a 19^th^ sickle haplotype was observed once in the ACB sample and a 20^th^ sickle haplotype was observed once in the Baganda (Table 3). The most common haplotype carried the ancestral allele at all 28 sites and accounted for 68.5% of all haplotypes. Thirteen of the sickle haplotypes in the Baganda, the one in the Zulu, and all four in the Qatari were identical to HAP1, the haplotype most commonly designated Central African Republic/Bantu (Table 3). Additionally, the autosomal fraction of African ancestry/patrilines/matrilines for the four Qatari carriers were 0.251/L1/L0a2a2a, 0.114/E1b1b1c*/H13c1, 0.974/E1b1a1a1f1a1/L3h1a2a1, and 0.078/NA/U6a2b1, indicating the presence of African ancestry in all four individuals. The three most common haplotypes (HAP1, HAP16, and HAP20) correlated primarily with the Central African Republic/Bantu, Benin, and Senegal designations, respectively.

**Table 3.**
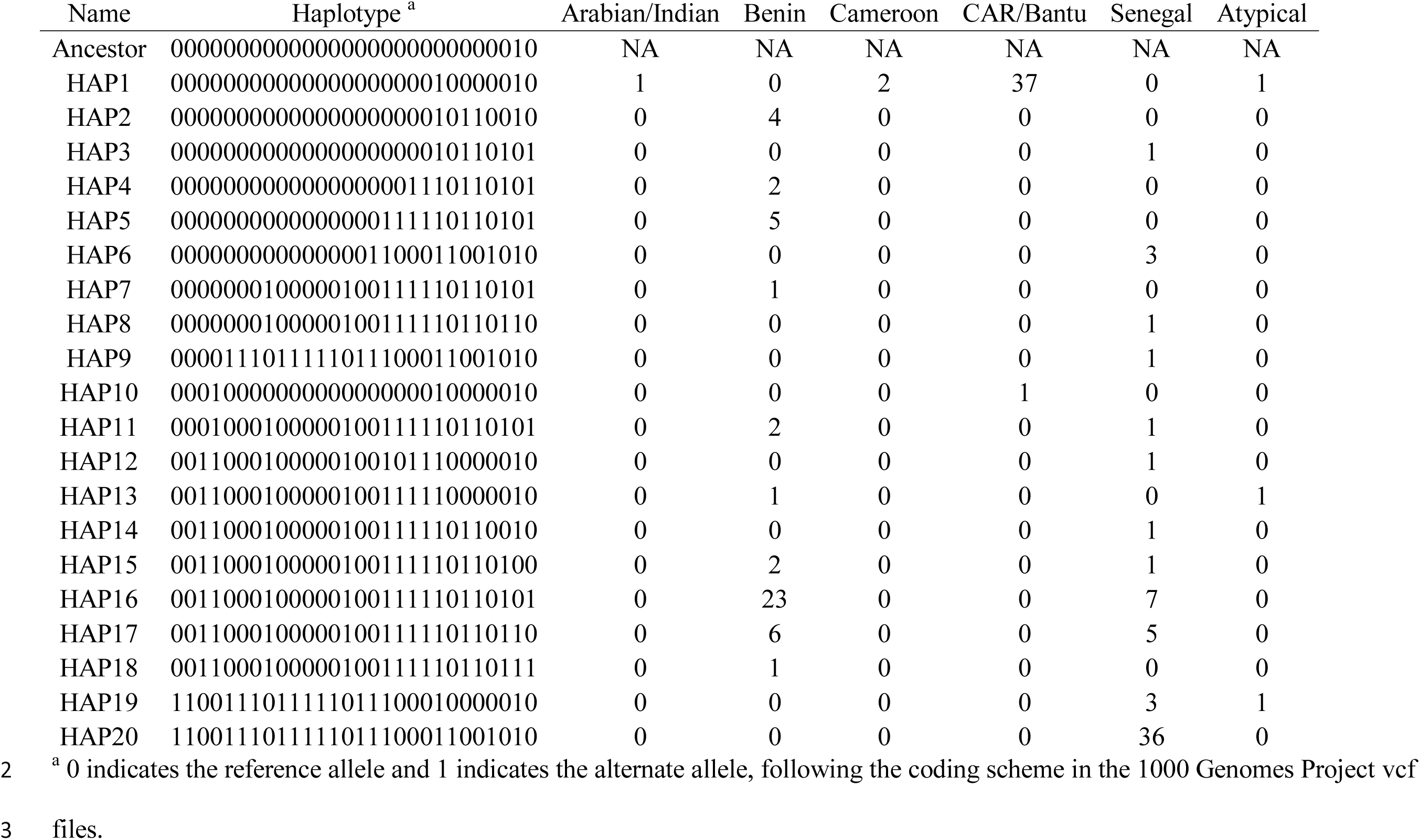
Distribution of sickle haplotypes under a sequence-based classification scheme.

### Bioinformatic annotation

Different haplotypes might be associated with different clinical phenotypes or disease severity.^40^ Using Ensembl and HaploReg version 4.1,^41^ we annotated each of the 27 sites in linkage disequilibrium with rs334 (Table S2). Possible modifiers include nine sites marked on histones as promoters or enhancers and three sites bound by proteins. In addition, rs1039215 is correlated with gene expression, most strongly with *HBG2* (Table S3). HAP1 (correlated with the Central African Republic/Bantu designation) and HAP16 (correlated with the Benin designation) differ by 13 sites, including rs73402608 (histone enhancer marks and bound protein) and rs1039215 (gene expression).

### Balancing selection

We modeled balancing selection assuming that the relative fitness of the β^A^/β^A^ homozygote was 1, the relative fitness of the β^S^/β^S^ homozygote was 0, and the relative fitness of the β^A^/β^S^ heterozygote was 1 + *s*. Based on the 504 continental Africans from the 1000 Genomes Project, we estimated that *s* = 0.158, in agreement with previous estimates.^14^ Next, we modeled random genetic drift plus balancing selection to estimate how many generations it would take for an equilibrium frequency of 12.0% to be reached, assuming a single initial copy and an effective population size *N*_*e*_ = 25542. We found that the mutant allele was lost 74.6% of the time and, conditional on reaching equilibrium, reached a frequency of 12.0% after an average of 87 (95% confidence interval [68,124]) generations. We stress that this value is not the age of the sickle mutation nor the age since the onset of balancing selection, but the time to reach a frequency of 12.0%. To determine the fate of the mutant allele in the absence of heterozygote advantage, we repeated the simulation assuming *s* = 0. We found that the mutant allele was lost after an average of 12 generations (95% confidence interval [1,92]), with a median of 2 generations.

### Phylogenetic network analysis

We used split decomposition analysis to infer the phylogeny of the 20 sickle haplotypes, rooted by the ancestral haplotype (Figure 1). The network revealed that the sickle mutation occurred once in the background of the ancestral haplotype and gave rise to HAP1, associated predominantly with the Central African Republic/Bantu designation. Two clusters were derived from this haplotype. One cluster contained HAP6, HAP9, HAP19, and HAP20, all associated with the Senegal designation. The other cluster contained haplotypes associated with both the Benin and Senegal designations.

**Figure 1.**
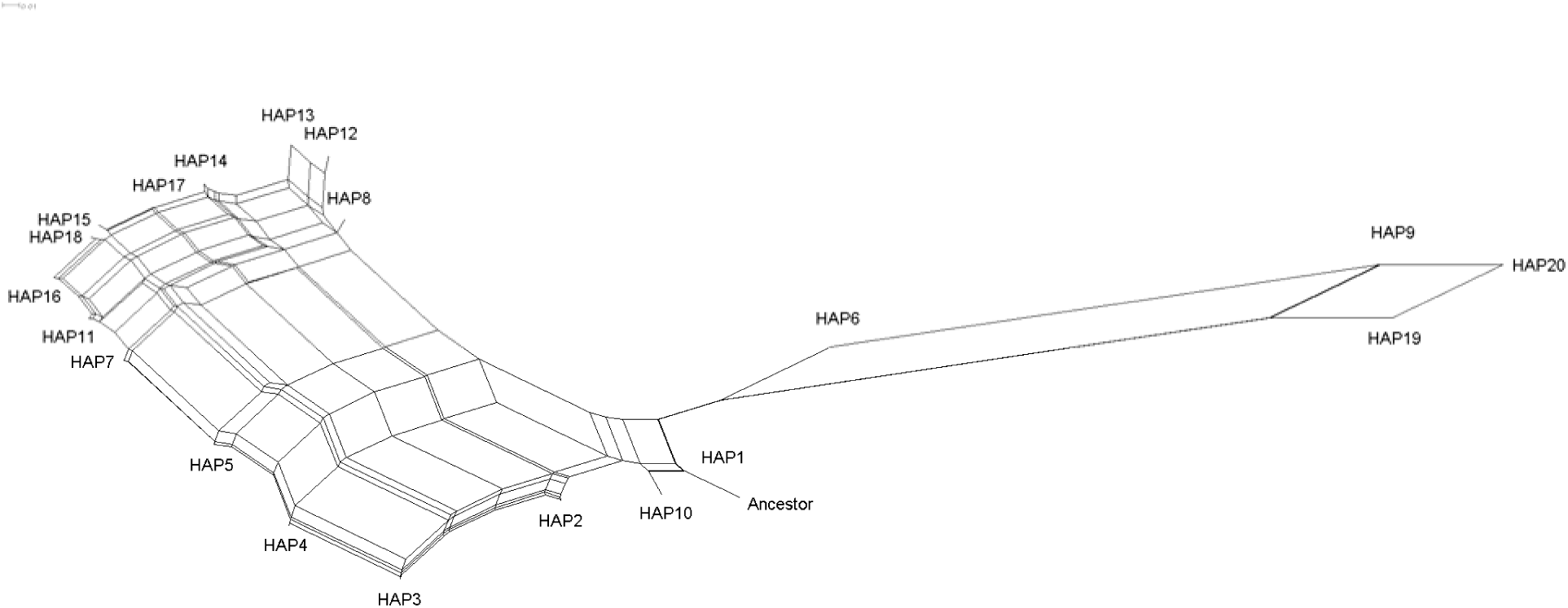
Split decomposition network of 20 distinct sickle-carrying haplotypes, rooted by the ancestral haplotype.

### The ancestral recombination graph

Using coalescent theory, we sampled the posterior distribution of the ancestral recombination graph using the 1,008 haplotypes, including 121 sickle haplotypes, from the 504 continental Africans from the 1000 Genomes Project. Conditional on this distribution, we estimated the age of the sickle mutation as 259 (95% confidence interval [123,395]) generations. Recombination rates of 1.7×10^−8^ recombinations/generation/site and larger yielded increasing numbers of incompatible sites.

## DISCUSSION

There are two models of the origins of the sickle allele. The multicentric model posits five independent occurrences of the same mutation in the last few thousand years. The unicentric model posits a single occurrence and an older age. We used whole genome sequence data to provide novel insight into this issue. Using a new haplotypic classification and phylogenetic network analysis, we found clear and consistent evidence for a single origin of the sickle mutation. After accounting for recombination, we estimated that the sickle mutation is 259 [123,395] generations old.

The earliest recorded cases of malaria were ~5,000 years ago.^2; 42; 43^ The first recorded cases of sickle were during the Hellenistic period, 2,130 years before present, in the Persian Gulf ^44^ and in 1670 AD in Ghana.^45^ Based on these limited data, it is historically plausible that malaria preceded the sickle mutation, consistent with our simulations of balancing selection showing that the sickle allele would have been lost almost immediately without a heterozygote advantage (assuming recessive lethality). The simulations of balancing selection also indicated that it took ~2,400 years for equilibrium to be reached. This time provides a lower bound on the age of the sickle mutation, since we do not know how long the equilibrium state has been maintained.

The Bantu Expansion started ~5,000 years ago.^46^ Our results imply that the sickle allele arose prior to the Bantu Expansion, consistent with the exclusive presence of the Central African Republic/Bantu haplotype in the Baganda and the Zulu. The Y chromosome haplogroups E1b1a1a1f and E1b1a1a1g, defined by L485 and U175 respectively, arose between 8,100 and 11,000 years before present. Our estimated age of the sickle mutation of ~7,300 years is consistent with a population in which these two sibling haplogroups co-circulated. Furthermore, our results place the origin of the sickle mutation in the middle of the Holocene Wet Phase or Neolithic Subpluvial, which lasted from ~7,500-7,000 BC to ~3,500-3,000 BC. This time was the most recent of the Green Sahara periods, during which the Sahara experienced wet and rainy conditions.^47^ Additionally, classical haplotypes in Western Arabia tend to have the Benin designation whereas those in Eastern Arabia tend to have the Arabian/Indian designation.^18^; ^21^ Although our sample includes only one predicted instance of the Arabian/Indian haplotype, the occurrence of this haplotype in the Luhya in Kenya and its clustering with the predominant haplotype found in Kenya and Uganda suggest an Eastern African waypoint. The statistical presence of African ancestry in all sickle carriers, combined with the statistical absence of Arabian or Indian ancestries in the five continental African samples in the 1000 Genomes Project,^32^ further supports an African origin of the sickle mutation.

We defined a new haplotypic classification based on phased sequence data. In contrast, the classical designations are based on restriction sites. By molecularly mapping restriction sites to the sequence data, we found that the restriction sites correlate poorly with rs334, such that none of the canonical sites was included in our sequence-based haplotypes. This result implies that the classical designations are not based on bona fide haplotypes. Our findings support a new classification based on three clusters. Notably, we found that the Senegal designation is substructured into two clusters, one of which shared with the Benin designation. This substructuring of haplotypes may have confounded previous assessments of clinical phenotype or disease severity.

## SUPPLEMENTAL DATA

Supplemental data include one figure and three tables.

## COMPETING INTERESTS

The authors declare no competing interests.

## ACKNOWLEDGEMENTS

We acknowledge the assistance of Neil Hanchard with the restriction site data. The contents of this publication are solely the responsibility of the authors and do not necessarily represent the official view of the National Institutes of Health. This research was supported by the Intramural Research Program of the Center for Research on Genomics and Global Health (CRGGH). The CRGGH is supported by the National Human Genome Research Institute, the National Institute of Diabetes and Digestive and Kidney Diseases, the Center for Information Technology, and the Office of the Director at the National Institutes of Health (1ZIAHG200362).

## WEB RESOURCES

Ensembl, http://www.ensembl.org/Homo_sapiens/Info/Index; E YTree, https://www.yfull.com/tree/E

